# Precision-Cut Liver Slices as an *ex vivo* model to evaluate antifibrotic therapies for liver fibrosis and cirrhosis

**DOI:** 10.1101/2023.10.30.564772

**Authors:** Yongtao Wang, Ben Leaker, Guoliang Qiao, Mozhdeh Sojoodi, Ibrahim Ragab Eissa, Eliana T. Epstein, Jonathan Eddy, Oizoshimoshiofu Dimowo, Georg M. Lauer, Raymond T. Chung, Motaz Qadan, Michael Lanuti, Bryan C. Fuchs, Kenneth K. Tanabe

## Abstract

**Background:** Precision-Cut Liver Slices (PCLS) are an *ex vivo* culture model developed to study hepatic drug metabolism. One of the main benefits of this model is that it retains the structure and cellular composition of the native liver. PCLS also represents a potential model system to study liver fibrosis in a setting that more closely approximates *in vivo* pathology than *in vitro* methods. The aim of this study was to assess whether responses to antifibrotic interventions can be detected and quantified with PCLS.

**Methods:** PCLS of 250 μm thickness were prepared from four different murine fibrotic liver models: choline-deficient, L-amino acid-defined, high-fat diet (CDAHFD), thioacetamide (TAA), diethylnitrosamine (DEN), and carbon tetrachloride (CCl_4_). PCLS were treated with 5 μM Erlotinib for 72 hours. Histology and gene expression were then compared with *in vivo* murine experiments and TGF-β1 activated hepatic stellate cells (HSCs). These types of PCLS characterization were also evaluated in PCLS from human cirrhotic liver.

**Results:** PCLS viability in culture was stable for 72 hours. Treatment of erlotinib, an EGFR inhibitor significantly inhibited the expression of profibrogenic genes *Il6*, *Col1a1* and *Timp1* in PCLS from CDAHFD-induced cirrhotic mice, and *Il6*, *Col1a1* and *Tgfb1* in PCLS from TAA-induced cirrhotic rats. Erlotinib treatment of PCLS from DEN-induced cirrhotic rats inhibited the expression of *Col1a1*, *Timp1*, *Tgfb1* and *Il6*, which was consistent with the impact of erlotinib on *Col1a1* and *Tgfb1* expression in *in vivo* DEN-induced cirrhosis. Erlotinib treatment of PCLS from CCl_4_-induced cirrhosis caused reduced expression of *Timp1*, *Col1a1* and *Tgfb1*, which was consistent with the effect of erlotinib in *in vivo* CCl_4_-induced cirrhosis. In addition, in HSCs at PCLS from normal mice, TGF-β1 treatment upregulated *Acta2* (*αSMA*), while treatment with erlotinib inhibited the expression of *Acta2*. Similar expression results were observed in TGF-β1 treated *in vitro* HSCs. Expression of MMPs and TIMPs, key regulators of fibrosis progression and regression, were also significantly altered under erlotinib treatment in PCLS. Expression changes under erlotinib treatment were also corroborated with PCLS from human cirrhosis samples.

**Conclusion:** The responses to antifibrotic interventions can be detected and quantified with PCLS at the gene expression level. The antifibrotic effects of erlotinib are consistent between PCLS models of murine cirrhosis and those observed *in vivo* and *in vitro*. Similar effects were also reproduced in PCLS derived from patients with cirrhosis. PCLS is an excellent model to assess antifibrotic therapies that is aligned with the principles of Replacement, Reduction and Refinement (3Rs).

## Introduction

Cirrhosis is the final stage of liver fibrosis caused by many forms of liver injury, such as chronic hepatitis B and C, chronic alcohol use disorder, and metabolic dysfunction-associated steatohepatitis (MASH, previously known as non-alcoholic steatohepatitis and NASH).^1, 2^ In 2015, cirrhosis affected 2.8 million people and resulted in 1.3 million deaths globally.^3, 4^ Cirrhosis also contributes to the vast majority of hepatocellular carcinoma (HCC) cases, which is the 6^th^ most common cancer worldwide, and the 2^nd^ leading cause of cancer-related death.^5^ Considering the lack of successful treatment options and poor prognosis for cirrhosis,^6^ new strategies and platforms to investigate antifibrotic therapies are urgently needed.

Many of the limitations of cellular and animal models of liver fibrosis can be overcome by using the relatively recently proposed precision cut liver slice (PCLS) model. This technique maintains thin slices of liver tissue in culture for several days, enabling drugs to be tested in an *ex vivo* setting that retains the architecture and cell populations of the liver. Compared to two-dimensional *in vitro* cell culture, PCLS better model the interactions and signal transduction pathways between different cell types.^7, 8^ The PCLS model also offers the benefit of reduced animal experiments and time compared to *in vivo* models, and is in line with the principles of Replacement, Reduction and Refinement.^9^

PCLS has been utilized to study various liver conditions, such as normal liver function,^10^ drug metabolism,^11^ drug-induced liver injury and toxicity,^12–14^ fatty liver,^15^ early stages of liver fibrosis,^8, 15–20^ and immunological responses.^21^ A study using PCLS derived from rats subjected to bile duct ligation supported PCLS’s ability for molecular antifibrotic evaluation with transcriptomic characterization.^22^ However, PCLS as a platform to evaluate antifibrotic therapies for cirrhosis, a more severe stage of liver disease, has not been reported.

Previous experiments from our lab have demonstrated the efficacy of a small-molecule EGF receptor (EGFR) inhibitor erlotinib as an antifibrotic agent in murine models of liver fibrosis.^23^ Erlotinib inhibited the activation of HSCs, stopped the progression of cirrhosis, and prevented subsequent development of HCC. Here, erlotinib was used as an example antifibrotic therapy in PCLS. The aim of this study was to assess whether responses to antifibrotic interventions can be detected and quantified with PCLS.

## Methods

### Chemicals

Stock solution of 50 mM erlotinib (Tarceva^TM^, Genentech) was prepared in DMSO.

### Animal models for PCLS

All animal experiments were approved by the MGH Institutional Animal Care and Use Committee (IACUC). PCLS were prepared from four established murine models of cirrhosis: choline-deficient, L-amino acid-defined, high-fat diet (CDAHFD), thioacetamide (TAA), diethylnitrosamine (DEN), and carbon tetrachloride (CCl_4_).

### DEN-induced cirrhosis rat model

The DEN model of cirrhosis was prepared as previously described.^23^ Male Wistar rats received weekly intraperitoneal (ip) injections of DEN (Sigma) at 50 mg/kg for 18 weeks.

### CCl_4_-induced cirrhosis mouse model

The CCl_4_ model of cirrhosis was prepared as previously described.^23^ Male C57Bl/6 mice were treated three times a week for 18 weeks with 0.1 ml of 40% CCl_4_ (Sigma) in olive oil by oral gavage.

### CDAHFD-induced cirrhosis mouse model

The CDAHFD model of cirrhosis was induced as previously described.^24^ Male C57Bl/6 mice received CDAHFD (L-amino acid diet with 60 kcal% fat with 0.1% methionine and no added choline. Research Diets Inc. A06071302) for 16 weeks.

### TAA-induced cirrhosis rat model

The TAA model of cirrhosis was induced as described.^25^ Male Wistar rats received ip injections with 200 mg/kg TAA (Sigma) twice a week for 12 weeks.

### PCLS as an *ex vivo* model of liver biology

PCLS from healthy or cirrhotic livers were performed as previously described.^7, 26, 27^ Tissue samples were glued to the mounting stage of a 7000smz-2 vibratome instrument (Campden Instruments Limited) and submersed in sterile Krebs Henseleit buffer (Sigma-Aldrich). Using 7550-1-C ceramic blades (Campden Instruments Limited), the tissue was cut into 250 μm slices at an advance speed of 0.1 mm/s, with 2.5 mm amplitude and 50Hz frequency. Slices from human and rat livers were trimmed to a uniform size with an 8 mm biopsy punch (Acuderm Inc.), and slices from mouse livers were trimmed with a 6 mm biopsy punch. PCLS were transferred into 8 μm-pore Transwell inserts and cultured in standard 6-well plates (Corning), with William’s E medium (Sigma-Aldrich) containing 2.0 g/L glucose, 10% fetal bovine serum (FBS, Gibco), 2 mM L-glutamine supplement (Gibco), 100 U/mL penicillin and 100 μg/mL streptomycin (Lonza), at 37°C in a humidified atmosphere of 5% CO_2_.

PCLS from cirrhotic animals were then treated with 5 μM erlotinib or vehicle control for 72 hours. PCLS from normal murine livers were treated with 10 ng/mL TGF-β1 for 24 hours, then treated with 10 ng/mL TGF-β1 and 5 μM erlotinib for an additional 72 hours.

### *In vivo* erlotinib experiments

#### DEN-induced cirrhosis rat model

As previously described,^23^ male Wistar rats received weekly ip injections of 50 mg/kg DEN for 18 weeks. Starting from week 8, rats received daily ip injections with either 2 mg/kg erlotinib or vehicle control (n = 8 per group). All rats were sacrificed at week 19 after a one-week washout period to eliminate the acute effects of DEN. The livers from this previous *in vivo* experiment were used in this current study.

#### CCl_4_-induced cirrhosis mouse model

As previously described,^23^ male A/J mice were treated three times a week for 18 weeks with 0.1 ml of 40% CCl_4_ in olive oil by oral gavage. Mice received daily ip injections of control or 2 mg/kg erlotinib from week 13 to 19 (n = 8 per group). Mice were sacrificed at week 19 after a one-week washout to eliminate acute the effects of CCl_4_. The livers from this previous *in vivo* experiment were used in this current study.

### Cell culture

Cell culture was performed as previously described.^28, 29^ Human HSCs, LX2 and TWNT4 cells were purchased from American Type Culture Collection (ATCC). Cells were grown in Dulbecco’s modified Eagle’s medium (DMEM, Cellgro) or Roswell Park Memorial Institute (RPMI) 1640 medium (HyClone) containing 15% FBS (Gibco), 1 mM sodium pyruvate (Gibco), 100 U/mL penicillin sodium and 100 μg/mL streptomycin sulfate (Lonza), at 37°C in a humidified atmosphere of 5% CO_2_.

### MTS

MTS assay was performed as previously described.^21^ Individual PCLS were placed in a 24-well plate with 400 μL William’s E medium and 80 μL MTS solution (Abcam) for each well. Plates were incubated at 37°C in standard culture conditions for 1 hour, then mixed briefly on a shaker. Supernatants were transferred to a 96-well plate and absorbance was measured using a plate reader (Molecular Devices) at OD = 490 nm.

### Histological Hematoxylin & Eosin (H&E) and Sirius red staining

Initially, PCLS were fixed in formalin for 3 days at room temperature then paraffin-embedded and sectioned into 7 μm slices for H&E and Sirius red staining with standard protocols.^30^ However, we found that PCLS from cirrhotic livers were fragile and easily degraded during sectioning and staining (**Fig. S3A**, left panel). The standard protocol was then modified. PCLS were fixed in formalin at 4°C overnight, embedded in paraffin, and then sectioned into 10 μm slices. The time for all steps were also reduced to half of the standard time. This modified protocol resulted in higher quality stains and retained the structure of the PCLS (**Fig. S3A**, right panel). Amount of collagen in Sirius red stained sections was scored according to the modified Ishak method^31^ as described in **Supplementary Table 1**. Collagen proportionate area was morphometrically quantified with image processing software (ImageJ, NIH).

### Quantitative RT-PCR

Quantitative RT-PCR was performed as previously described.^32, 33^ Total RNA was isolated from rat liver tissues using TRIzol (Invitrogen) and subsequently treated with DNase I (Promega) according to the manufacturer’s instructions. 1 μg of total RNA from each sample was used to synthesize cDNA (SuperScript III First-Strand Synthesis SuperMix for qRT-PCR, Invitrogen). Quantitative real-time PCR was performed using the 7900HT Fast Real-Time PCR System (Thermo fisher) with commercial TaqMan primers (Thermo fisher) as shown in **Supplementary Table 2**.

### Human samples

All human samples were harvested in accordance with the protocols approved by the Mass General Brigham Institutional Review Boards (IRBs).

### Statistical analysis

All values were expressed as mean ± S.E.M.. Two-tailed Student’s t tests were performed to compare data between control and one experimental group, and One-way ANOVA followed by post-hoc Tukey tests with 2-tailed distribution were performed to analyze data among groups of 3 or more. Graphs were prepared with GraphPad Prism v6.0c software. All experiments were independently repeated three times. Significance is represented by \**p* < 0.05, \*\**p* < 0.01, \*\*\**p* < 0.001 versus control.

## Results

### Stable viability of PCLS

PCLS with 250 μm thickness were prepared and further cultured with treatment (**Fig. 1A, S1**). To ensure minimal tissue degradation *ex vivo,* the period of stable viability was determined. MTS assays showed the viability of PCLS from mouse CDAHFD-induced, rat DEN-induced, rat TAA-induced, and mouse CCl_4_-induced established cirrhosis were not significantly changed after 72 hours in culture (**Fig. 1B-1E, S2A, S2B**). This timepoint was used for the remaining experiments with cirrhotic PCLS. The DEN model was also used to investigate the lifespan of cirrhotic PCLS. The nominal decrease in viability after 5 days was not statistically significant (**Fig. 1C**). The CDAHFD and DEN models were further used to determine whether erlotinib had an impact on PCLS viability. There was no difference between vehicle and erlotinib treated slices with either model at 72 hours (**Fig. S2C, S2D**).

**Figure 1.**
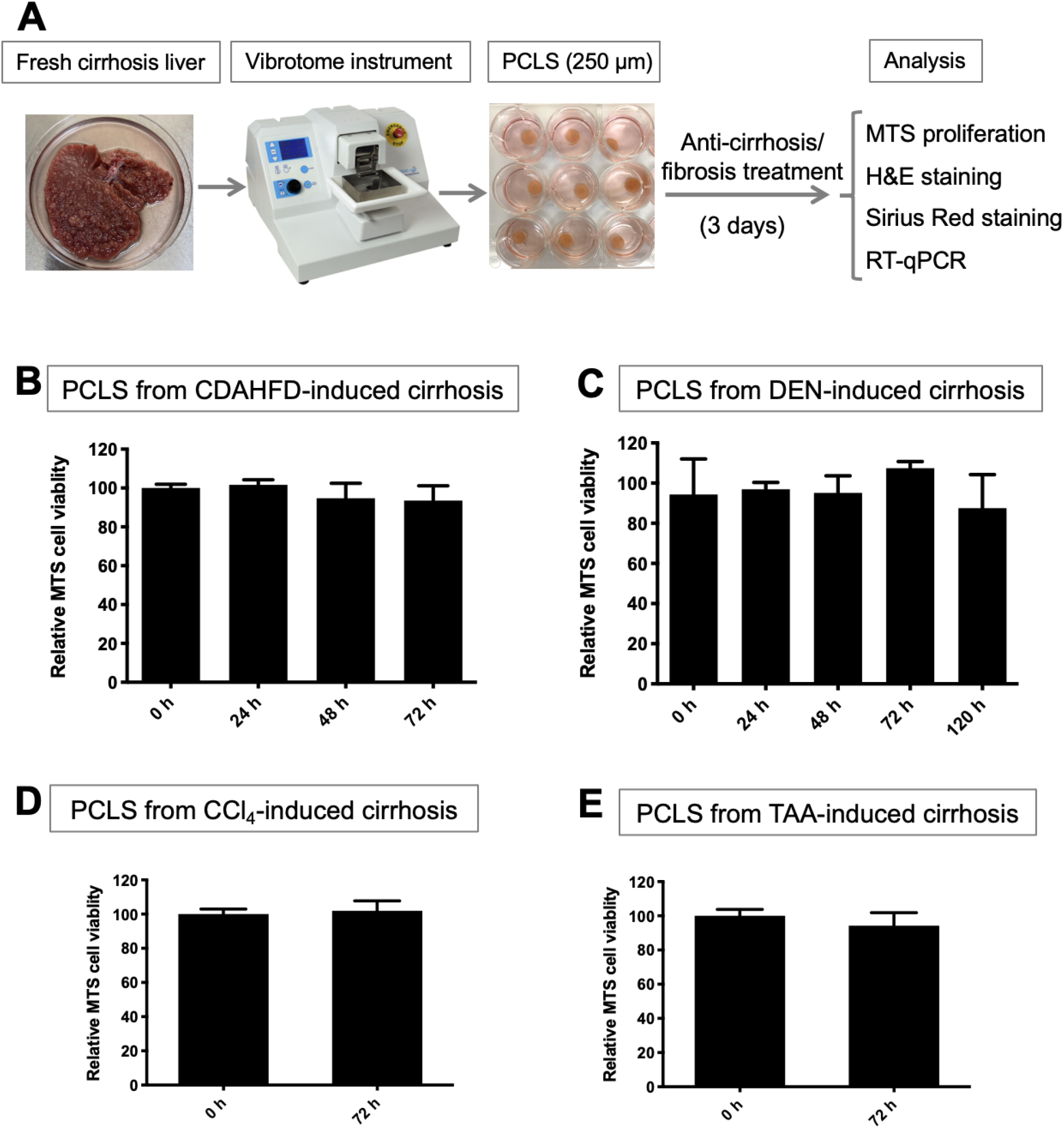
Evaluation of viability in PCLS. (A) Workflow for PCLS preparation and treatment. Viability analyzed with MTS assays for PCLS from (B) CDAHFD, (C) DEN, (D) CCl_4_ and (E) TAA-induced cirrhosis rats. Experiments were independently repeated three times (n = 3). All values were expressed as the mean ± S.E.M with two-tailed Student’s *t* tests or one-way ANOVA analysis.

### Antifibrotic treatment of PCLS from CDAHFD-induced mouse cirrhosis and TAA-induced rat cirrhosis

To first assess whether responses to antifibrotic interventions can be detected and quantified with PCLS, PCLS from CDAHFD-induced and TAA-induced cirrhosis were utilized. CDAHFD-induced cirrhosis^30^ was confirmed with Sirius red staining (**Fig. S4A)**, and PCLS from this CDAHFD-induced cirrhosis were prepared. Erlotinib treatment of PCLS slices for 72 hours significantly suppressed the expression of the profibrogenic genes *Il6*, *Col1a1* and *Timp1* (**Fig. 2A**), and suppressed *Acta2* expression with marginal significance (*p* = 0.0777). No significant effect was observed on the expression of *Tgfb1*. This short-term exposure of PCLS slices to erlotinib did not significantly reduce the amount of collagen measured with Sirius red staining (**Fig. 2B, 2C**). No change was evident on H&E stained morphology (**Fig. S5A**).

**Figure 2.**
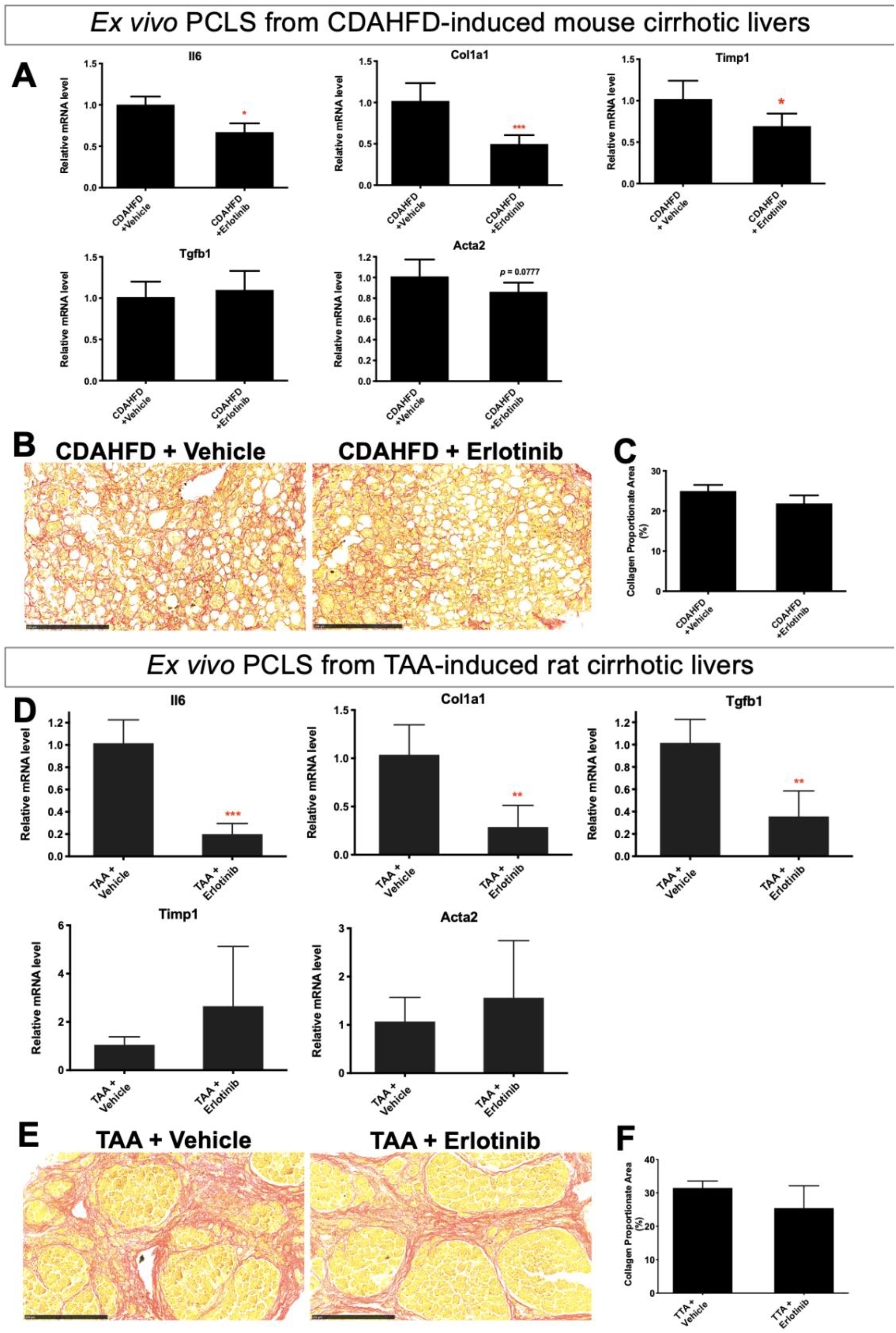
Antifibrotic evaluation in PCLS from CDAHFD-induced mouse cirrhosis and TAA-induced rat cirrhosis. (A) Quantitative RT-PCR analysis of profibrogenic genes. Amount of collagen (B) measured with Sirius red staining and (C) quantified with collagen proportionate area analysis in PCLS from CDAHFD-induced mouse cirrhosis after 5 μM erlotinib treatment for 72 hours. (D) Quantitative RT-PCR analysis of profibrogenic genes. Amount of collagen (E) measured with Sirius red staining and (F) quantified in PCLS from TAA-induced rat cirrhosis after 5 μM erlotinib treatment for 72 hours. Experiments were independently repeated three times (n = 4). All values were expressed as the mean ± S.E.M with two-tailed Student’s *t* tests. Significance is represented by \**p* < 0.05, \*\**p* < 0.01, \*\*\**p* < 0.001.

Cirrhosis was induced by TAA^34^ and confirmed with Sirius staining (**Fig. S4B**). PCLS were prepared from these TAA-induced cirrhotic livers. Erlotinib treatment of PCLS slices for 72 hours significantly inhibited the expression of the profibrogenic genes *Il6*, *Tgfb1* and *Col1a1* in PCLS (**Fig. 2D**). No significant change in expression of *Timp1* and *Acta2* was observed. This short-term exposure to erlotinib did not significantly reduce the amount of collagen (**Fig. 2E, 2F**). No change was evident on H&E staining (**Fig. S5B**).

Taken together with the CDAHFD and TAA data above, acute responses to antifibrotic interventions can be detected and quantified with PCLS on the molecular gene expression level.

### Antifibrotic treatment in PCLS from DEN-induced rat cirrhosis compared to *in vivo* DEN cirrhosis model

DEN-induced cirrhosis^23, 35, 36, 37, 38^ was confirmed with Sirius staining (**Fig. S4C**), and PCLS from this DEN-induced cirrhosis were then prepared. Exposure of PCLS slices to erlotinib for 72 hours significantly suppressed the expression of the profibrogenic genes *Col1a1*, *Tgfb1*, *Il6* and *Timp1* (**Fig. 3A**). No significant difference in the amount of collagen (**Fig. 3B, 3C**) and H&E staining (**Fig. S5C**) were observed after erlotinib treatment of PCLS slices for 72 hours.

**Figure 3.**
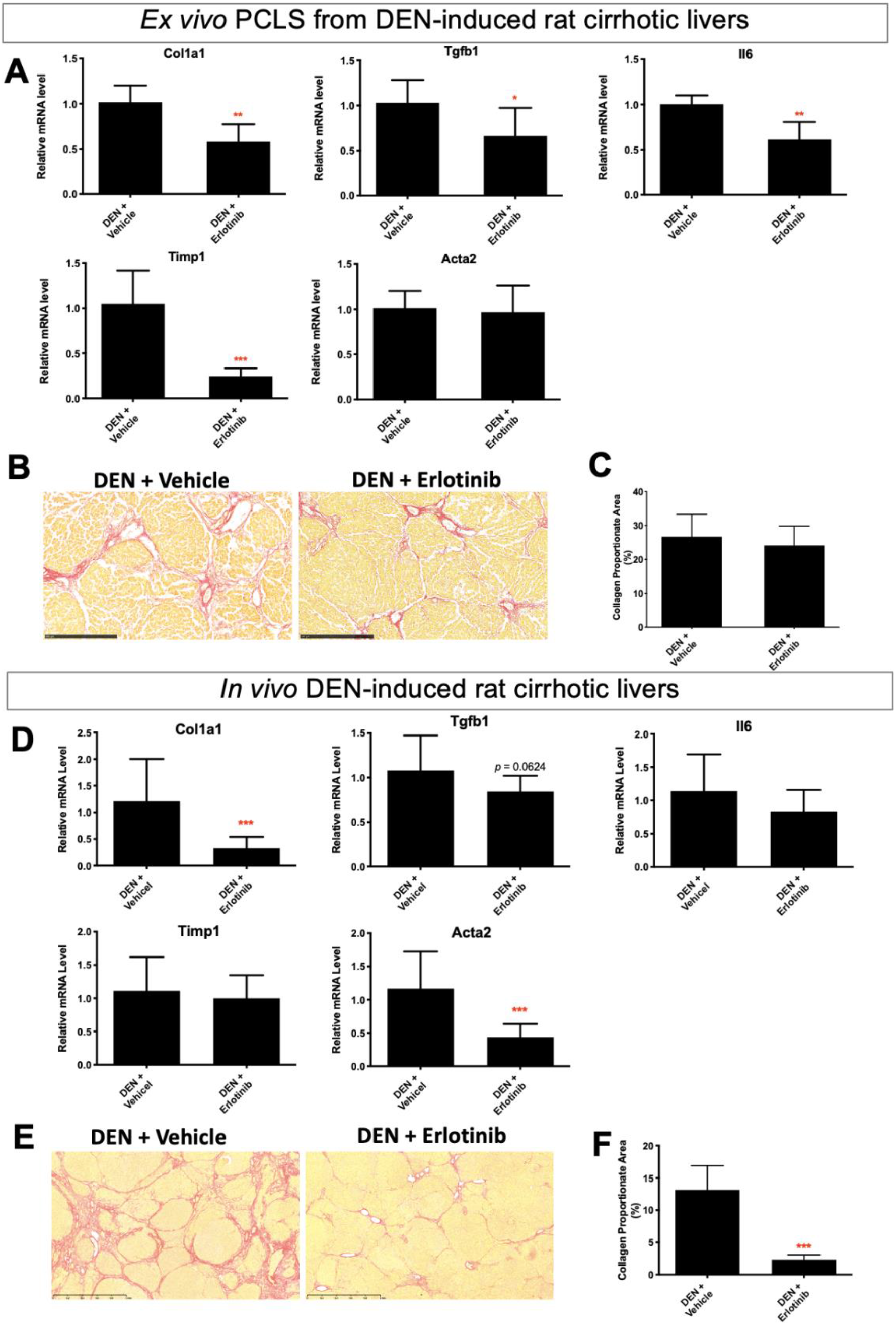
Comparison of antifibrotic therapy in *ex vivo* PCLS from DEN-induced *rat* cirrhosis with *in vivo* DEN cirrhotic model. For *ex vivo* PCLS analysis, (A) quantitative RT-PCR analysis of profibrogenic genes. Amount of collagen (B) measured with Sirius red staining and (C) quantified with collagen proportionate area analysis in PCLS after 5 μM erlotinib treatment for 72 hours. Experiments were independently repeated three times (n = 4). For *in vivo* analysis, (D) quantitative RT-PCR analysis of profibrogenic genes. Amount of collagen (E) measured with Sirius red staining and (F) quantified after daily ip injections of 2 mg/kg erlotinib for 10 weeks. All values were expressed as the mean ± S.E.M with two-tailed Student’s *t* tests. Significance is represented by \**p* < 0.05, \*\**p* < 0.01, \*\*\**p* < 0.001.

To compare the effect of erlotinib between *ex vivo* cirrhotic PCLS and an *in vivo* cirrhosis model, we analyzed livers obtained from DEN-induced cirrhotic rats that had been treated with erlotinib for 10 weeks (**Fig. S5E**). As observed in the short-term PCLS model, longer-term treatment rats with DEN-induced cirrhosis resulted in significant reduction in *Col1a1* and *Acta2* (**Fig. 3D**), and marginally reduced the expression of *Tgfb1* (*p* = 0.0624). No effect on the expression of *Il6* and *Timp1* was observed. As previously published, 10 weeks of erlotinib treatment in these rats also significantly reduced the amount of collagen (**Fig. 3E, 3F**).

### Antifibrotic treatment in PCLS from CCl_4_-induced mouse cirrhosis compared to *in vivo* CCl_4_ cirrhosis model

CCl_4_-induced cirrhosis^23^ was confirmed with Sirius red staining (**Fig. S4D**) and PCLS from this CCl_4_-induced cirrhosis were then prepared. Short-term exposure of PCLS slicers to erlotinib significantly suppressed the expression of the profibrogenic genes *Timp1*, *Col1a1* and *Tgfb1* (**Fig. 4A**), but did not affect the expression of *Il6* and *Acta2*. No significant difference in the amount of collagen **(Fig. 4B, 4C**) and H&E staining (**Fig. S5D**) were observed after erlotinib treatment of the PCLS for 72 hours.

**Figure 4.**
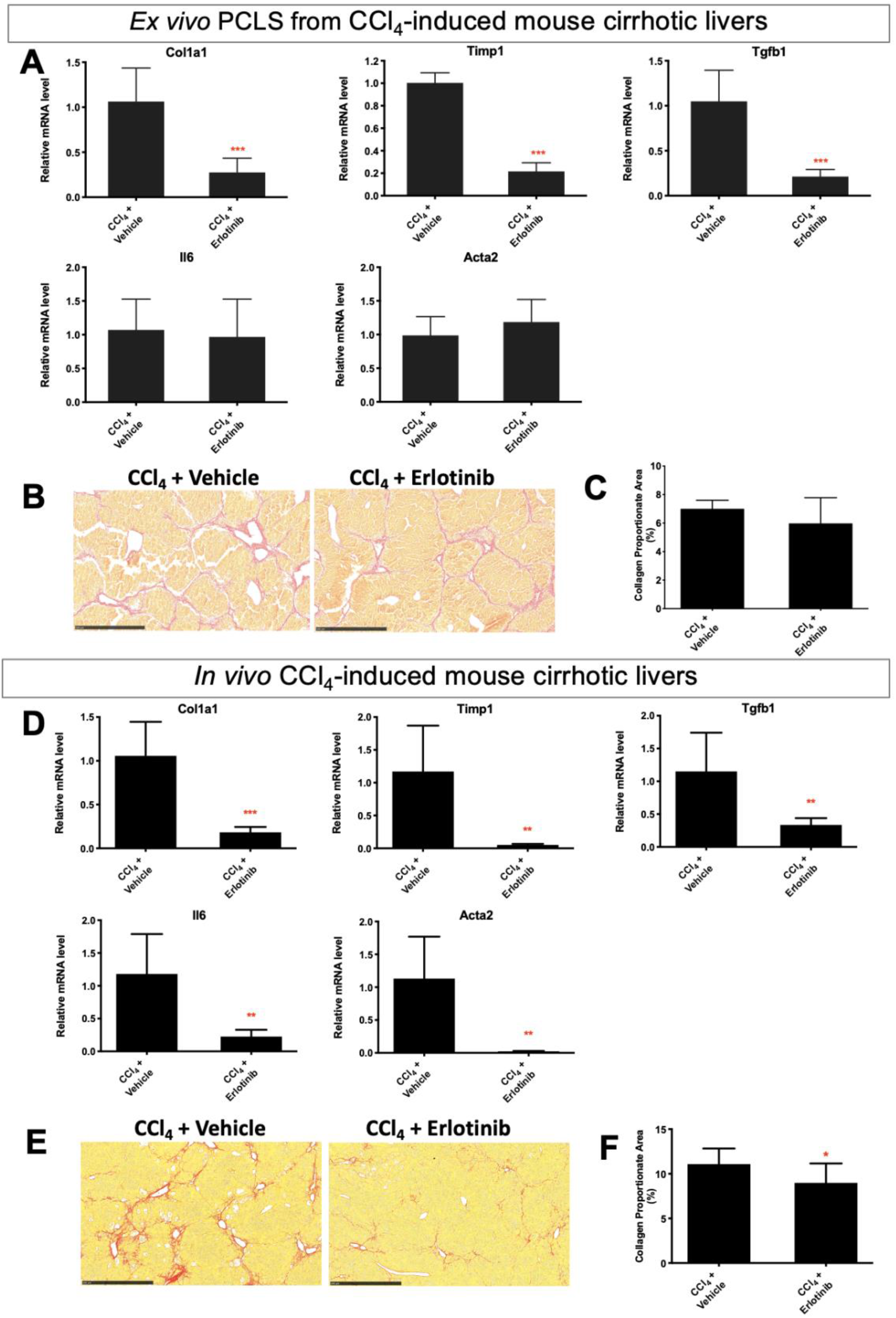
Comparison of antifibrotic therapy in *ex vivo* PCLS from CCl_4_-induced mouse cirrhosis with *in vivo* CCl_4_ cirrhotic model. For *ex vivo* PCLS analysis, (A) quantitative RT-PCR analysis of profibrogenic genes. Amount of collagen (B) measured with Sirius red staining and (C) quantified with collagen proportionate area analysis in PCLS after 5 μM erlotinib treatment for 72 hours. Experiments were independently repeated three times (n = 4). For *in vivo* analysis, (D) quantitative RT-PCR analysis of profibrogenic genes. Amount of collagen (E) measured with Sirius red staining and (F) quantified after daily ip injections of 2 mg/kg erlotinib for 6 weeks. All values were expressed as the mean ± S.E.M with two-tailed Student’s *t* tests. Significance is represented by \**p* < 0.05, \*\*\**p* < 0.001.

To compare the effect of erlotinib between *ex vivo* cirrhotic PCLS and an *in vivo* cirrhosis model, we analyzed liver obtained from CCl_4_-induced cirrhotic mice that had been treated with erlotinib for 6 weeks (**Fig. S5F**). Six weeks of erlotinib treatment in these mice significantly suppressed the expression of *Timp1*, *Col1a1*, *Tgfb1*, *Il6* and *Acta2* (**Fig. 4D**), and reduced the amount of collagen (**Fig. 4E, 4F**).

Taken together with the DEN and CCl_4_ data above, these results demonstrate that erlotinib induces acute effects in PCLS that are similar to those observed *in vivo* in these two murine models of cirrhosis.

### Antifibrotic treatment in HSCs from PCLS compared to *in vitro* HSCs

In PCLS from normal rats (**Fig. S5G**), HSCs which expressing *Acta2* were only a small minority of cells. TGF-β1 treatment for 72 hours induced the expression of *Acta2* in HSCs at PCLS, while treatment with erlotinib significantly inhibited the expression of *Acta2* (**Fig. 5A**). In stellate cell lines LX2 (**Fig. 5B**) and TWNT4 (**Fig. 5C**), TGF-β1 significantly induced the expression of *ACTA2*, while treatment with erlotinib for 72 hours suppressed *ACTA2* expression. Erlotinib treatment for 72 hours also reduced TGF-β1-induced collagen deposition (**Fig. 5D**) on the morphological level. These results demonstrate that erlotinib induces similar effects in HSCs in PCLS and *in vitro* HSCs.

**Figure 5.**
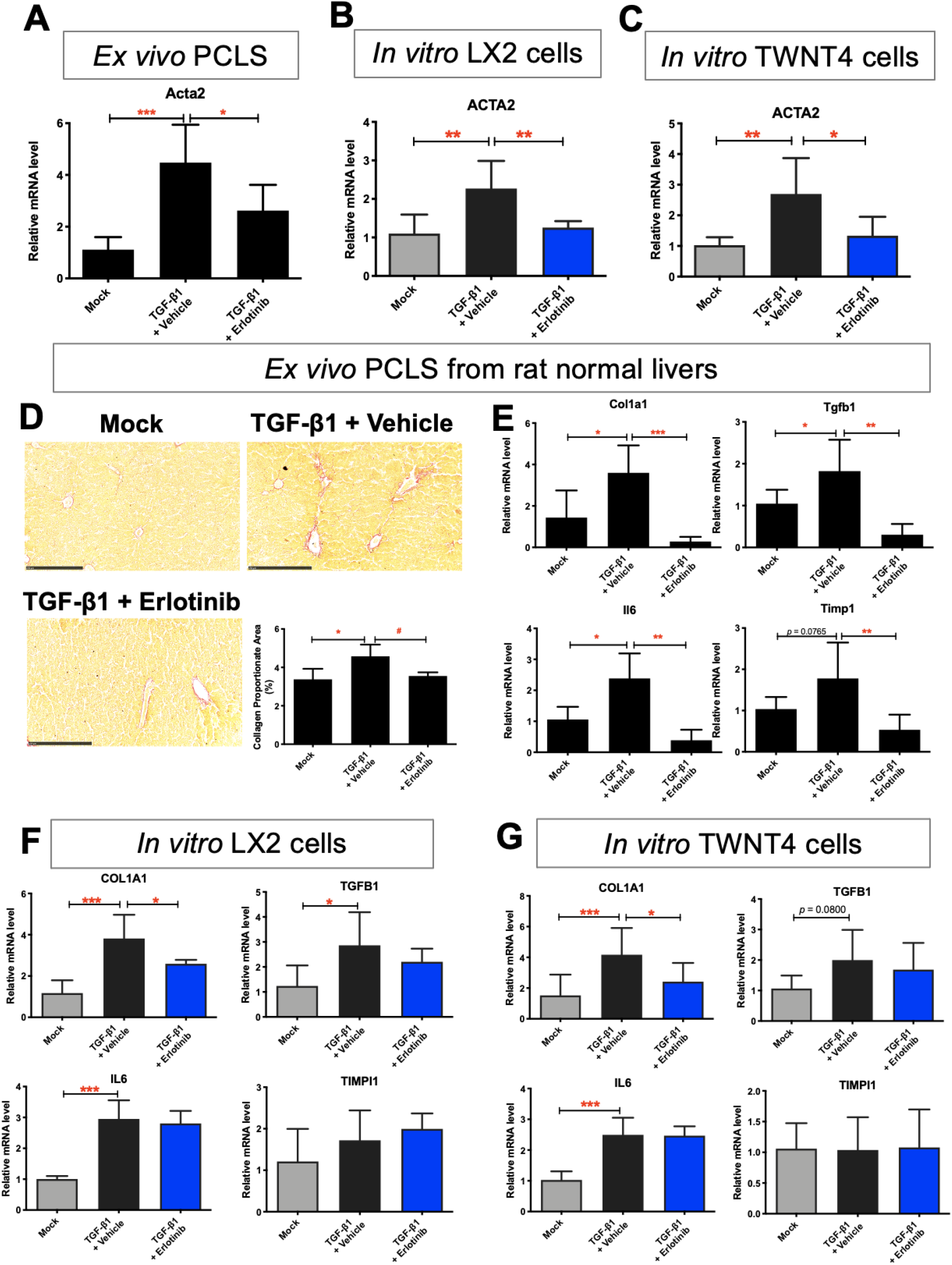
Comparison of antifibrotic evaluation in *ex vivo* HSCs from PCLS with *in vitro* HSCs. Quantitative RT-PCR analysis of *Acta2* mRNA from (A) PCLS from normal rats, (B) LX2 and (C) TWNT4 cells. PCLS or HSCs were activated with 10 ng/mL TGF-β1 for 24 hours, and further treated with 5 μM erlotinib for 72 hours. (D) Amount of collagen measured with Sirius red staining and quantified with collagen proportionate area analysis in PCLS from normal rats. Quantitative RT-PCR analysis of *Il6*, *Tgfb1*, *Timp1* and *Col1a1* mRNA from (E) PCLS from normal rats, (F) LX2 and (G) TWNT4 cells. These experiments were repeated three times (n = 4). All values were expressed as the mean ± S.E.M with one-way ANOVA analysis. Significance is represented by \**p* < 0.05, \*\**p* < 0.01, \*\*\**p* < 0.001.

Besides, TGF-β1 treatment induced the expression of other profibrogenic genes *Il6*, *Tgfb1 Timp1* and *Col1a1* in PCLS, while treatment with erlotinib significantly inhibited the expression of these genes (**Fig. 5E**). In stellate cell lines LX2 (**Fig. 5F**) and TWNT4 (**Fig. 5G**), TGF-β1 significantly induced the expression of *COL1A1*, while treatment with erlotinib suppressed *COL1A* expression. However, *TGFB1* and *IL6* did not respond to erlotinib treatment, and *TIMP1* did not respond to TGF-β1 and erlotinib treatment on these cells.

### Effect of antifibrotic therapy on MMPs and TIMPs expression in cirrhotic PCLS

To explore other potential mechanisms of antifibrotic therapy of erlotinib in cirrhotic PCLS, we assessed expression of other regulatory pathways for fibrosis generation and degradation. In livers from CDAHFD-induced cirrhotic mice, exposure of PCLS slices to erlotinib for 72 hours significantly increased the expression of *Mmp2*, *Mmp3* and *Mmp8* (**Fig. 6A**), indicating activation of antifibrotic pathways. Erlotinib treatment also significantly decreased *Mmp9*, *Mmp13* (**Fig. 6B**), and *Timp1* (**Fig. 6C**) expression, reflecting suppression of profibrotic pathways. Decreased *Timp1* expression was also observed in PCLS from CDAHFD (**Fig. 2A**), DEN (**Fig. 3A**), and CCl_4_-induced (**Fig. 4A**) cirrhotic livers.

**Figure 6.**
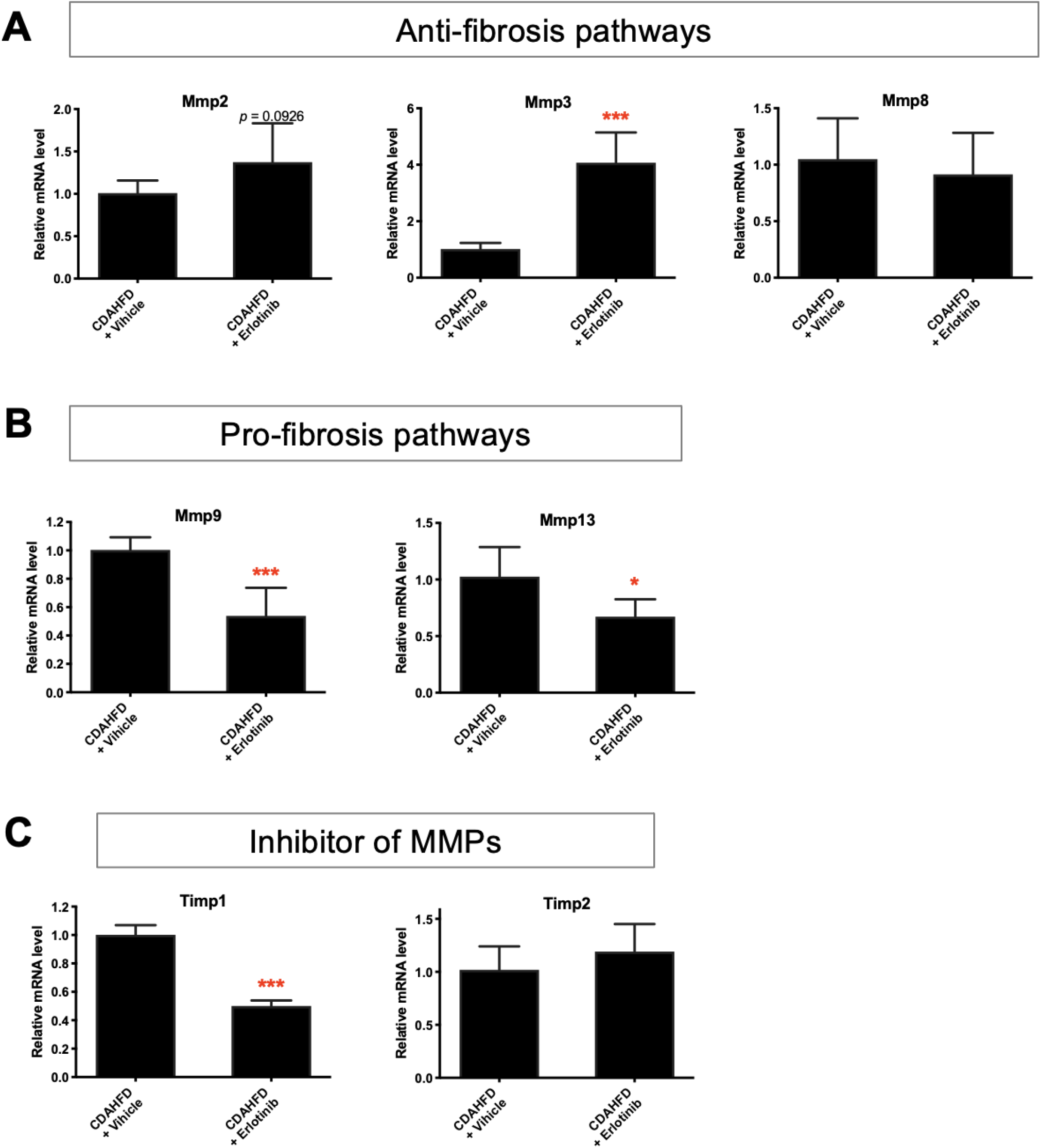
Regulatory pathways of the antifibrotic evaluation of erlotinib in cirrhotic PCLS. Quantitative RT-PCR analysis of the mRNA expression on (A) antifibrotic pathways, (B) profibrotic pathways, and (C) inhibitory pathways of MMPs in PCLS from CDAHFD-induced cirrhosis mice after erlotinib treatment for 72 hours. These experiments were repeated three times (n = 4). All values were expressed as the mean ± S.E.M with two-tailed Student’s *t* tests. Significance is represented by \**p* < 0.05, \*\*\**p* < 0.001.

### Antifibrotic evaluation in human cirrhotic PCLS

The PCLS from diet-induced murine cirrhotic model and cirrhotic patients were utilized to assess whether the types of PCLS characterizations in murine cirrhotic PCLS were also observed in PCLS from human cirrhosis. As described above, PCLS from CDAHFD-induced murine cirrhosis was used to assess the effects of 72 hours exposure to erlotinib (**Fig. 2A-C**). We next tested the effects of 72 hours of erlotinib on PCLS from human liver cirrhosis (**Fig. S6**). Erlotinib treatment of PCLS did not change the liver morphology (**Fig. S5H**), but significantly inhibited the expression of the profibrogenic genes *COL1A1*, *TIMP1*, *IL6* and *TNFA* (**Fig. 7A**). No significant effect was observed on the expression of *Tgfb1* and *Acta2*. No significant difference in collagen accumulation was observed after erlotinib treatment (**Fig. 7B, 7C**). Analysis using PCLS demonstrated that the effects of erlotinib as an antifibrotic therapy were similar in murine and human cirrhosis samples.

**Figure 7.**
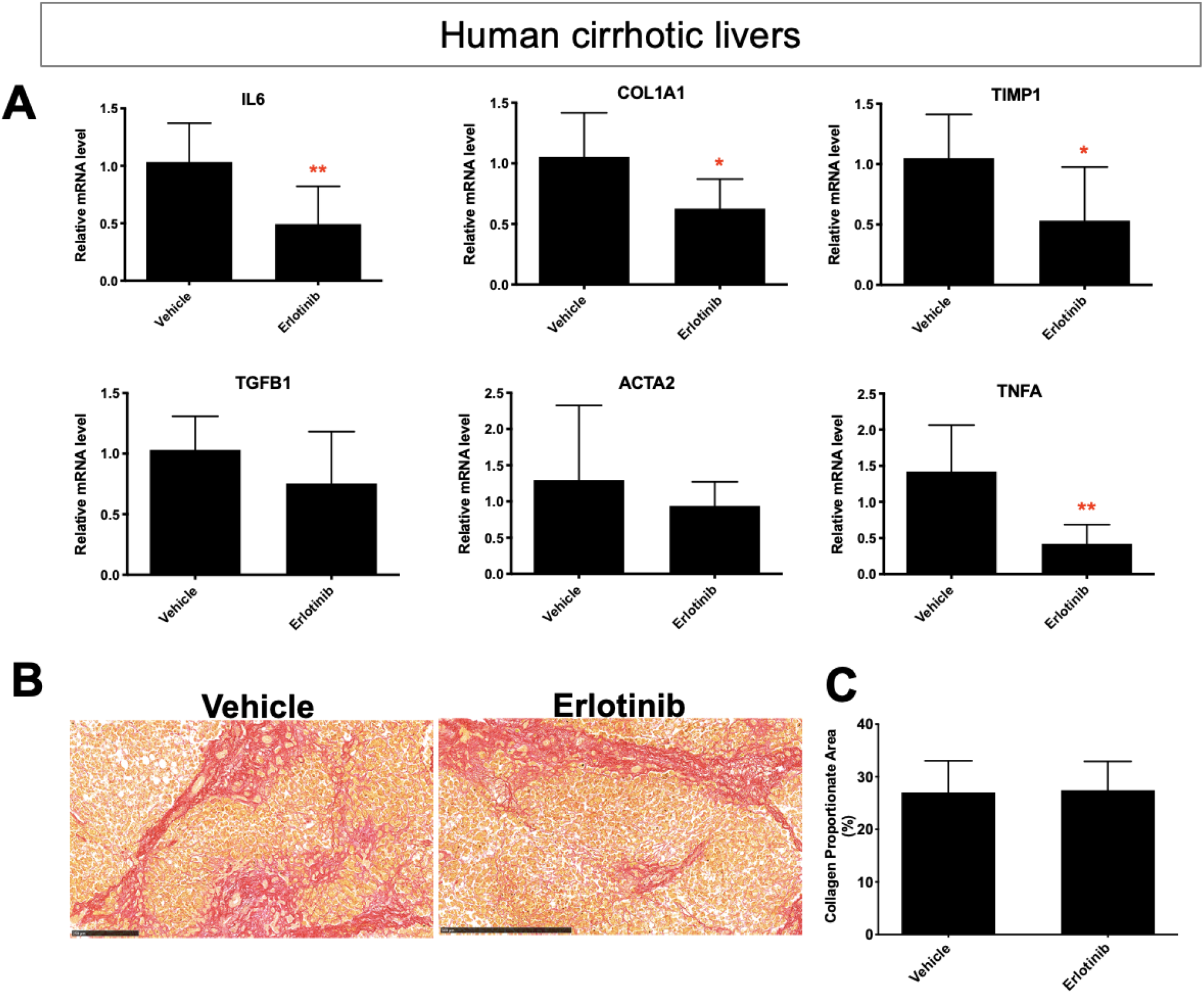
The Antifibrotic evaluation in human cirrhotic PCLS. (A) Quantitative RT-PCR analysis of mRNA expression of profibrogenic genes. Amount of collagen (B) measured with Sirius red staining and (C) quantified in PCLS from human cirrhosis treated with 5 μM erlotinib for 72 hours. These experiments were repeated three times (n = 4). All values were expressed as the mean ± S.E.M with two-tailed Student’s *t* tests. Significance is represented by \*\**p* < 0.01.

## Discussion

New strategies to investigate antifibrotic therapies for cirrhosis are urgently needed. Compared with two-dimensional *in vitro* cell lines, PCLS are a better representation of the *in vivo* environment because they retain the structure and cellular composition.^7, 8^ It therefore has the capacity to better capture the complex multi-cellular pathways involved in liver injury and the progression of cirrhosis. Compared to *in vivo* models, PCLS offer faster readouts, reduced animal numbers, and the ability to run multiple tests on highly similar samples by using serial sections.

In this study, we demonstrated that PCLS is a reliable *ex vivo* model to evaluate antifibrotic therapies across four established murine models of (CDAHFD, TAA, DEN, and CCl_4_-induced) cirrhosis, using erlotinib as an example drug. Expression analysis of PCLS after erlotinib treatment showed suppression in a variety of profibrogenic genes. We demonstrated the responses to antifibrotic interventions can be detected and quantified with PCLS at the molecular level. As expected, morphology of PCLS after erlotinib treatment did not significantly change. This is likely because the short time in culture (72 hours) does not provide enough time for significant remodeling to occur.^7^ Our previous i*n vivo* experiments showing reduced collagen accumulation in DEN-induced cirrhosis in rats or CCl_4_-induced cirrhosis in mice used erlotinib treatment for longer-term weeks.^23^ Our current study also confirmed that the viability of PCLS from cirrhotic liver stabilized during 72 hours in culture.

We directly compared PCLS with *in vivo* experiments using the same method of inducing cirrhosis, and showed that many of the acute responses to antifibrotic therapies in PCLS were consistent with *in vivo* results. Erlotinib treatment of PCLS from DEN-induced cirrhotic rats suppressed the expression of partial profibrogenic genes, which was consistent with the impact of erlotinib on these genes *in vivo*. Similar results were demonstrated with the CCl_4_ mouse model.

HSCs expressing *Acta2* are a small minority population within PCLS. We also compared HSCs at PCLS to common *in vitro* HSCs. TGF-β1 activated LX2 and TWNT4 cell lines are well established models to evaluate antifibrotic therapies *in vitro*.^36, 39^ Here we showed erlotinib treatment significantly suppressed expression of the TGF-β1 induced profibrogenic gene *Acta2* in HSCs on normal liver PCLS. Erlotinib also significantly inhibited the expression of TGF-β1 activated *ACTA2* in LX2 and TWNT4 cells. As other profibrogenic genes *Il6*, *Tgfb1*, *Timp1* and *Col1a1* are secreted by hepatocytes, HSCs and other types of liver cells,^40–43^ the decreased expression of these genes in PCLS after erlotinib treatment might come from all these liver cells. However, erlotinib only significantly inhibited the expression of TGF-β1 activated *COL1A1* in LX2 and TWNT4 cells. *IL6*, *TIMP1* and *TGFB1* were not reduced after erlotinib treatment, which might also indicate a lower sensitivity for cell lines to respond to certain antifibrotic therapies.

We then investigated other potential mechanisms contributing to these molecular observations of fibrogenesis in PCLS. Matrix metalloproteinases (MMPs) are a family of enzymes that regulate the degradation of extracellular matrix proteins. MMP13 induces inflammation, cytokines and activation of myofibroblast to acquire the profibrotic effect. On the contrary, MMP2 and MMP8 induce clearance of collagen, and MMP3 inhibits activation of myofibroblast to foster the anti-fibrosis effect. Besides, lack of MMP9 results in reduced fibrosis. Tissue inhibitors of metalloproteinases (TIMPs) inhibit the MMPs.^44^ Regulatory pathways on the fibrosis generation and degradation^22, 44^ were significantly altered to exert the antifibrotic therapy of erlotinib in PCLS. After erlotinib treatment, antifibrotic pathways including MMP2, MMP3 and MMP8 were enhanced in PCLS from CDAHFD-induced cirrhosis, while profibrotic pathways including MMP9 and MMP13 were hampered. Besides, TIMP1 as the inhibitor of MMPs, was reduced after erlotinib treatment.

We then confirmed the antifibrotic effects of erlotinib using PCLS from human cirrhosis samples. The type of PCLS characterizations in murine cirrhotic PCLS were also observed in PCLS from human cirrhosis, which addressed the prospect to further assess the effect of erlotinib in *in vivo* human cirrhotic liver. The majority of PCLS research to date has focused on murine samples^8, 10, 11, 13–19^ and only a small fraction of studies used human tissue.^12, 20^ PCLS from human samples may better predict the *in vivo* response to an antifibrotic therapy in humans, and also opens up the possibility of using a liver biopsy sample to test an individual patient’s response to a variety of drugs. Human PCLS may also be used as an experimental platform to expedite basic research and drug development, by directly testing the effects of drugs in clinical cirrhosis samples rather than animal models.

In summary, the responses to antifibrotic interventions can be detected and quantified with PCLS on the molecular level. PCLS accurately capture the changes in expression that occur *in vitro* and *in vivo* during treatment with an antifibrotic therapy. These types of PCLS characterizations were also observed in PCLS from human cirrhosis. PCLS reduces animal numbers, in alignment with the principles of Replacement, Reduction and Refinement^9^ by enabling many slices to collected from one animal. PCLS is a promising platform for the future development of antifibrotic therapies for cirrhosis.

## Acknowledgements

This study has been supported by R01DK104956, NIH. We acknowledge the kind help from Dr. Alan C. Mullen, who previously worked at Liver Center, Division of Gastroenterology, Massachusetts General Hospital and Harvard Medical School, and currently works at Division of Gastroenterology, University of Massachusetts Chan Medical School and Broad Institute, to get access to the human liver samples.

## Author contributions

KKT and YW designed the experiments. YW, BL and GQ conducted the experiments and performed data analysis. YW wrote the manuscript, and KKT and BL improved the manuscript. MS, IRE, KK, SCB, ETE, JE, GML, RTC, MQ, ML, BCF and RTC contributed to new reagents or analytical tools.

---

**Supplementary Figure 1.**
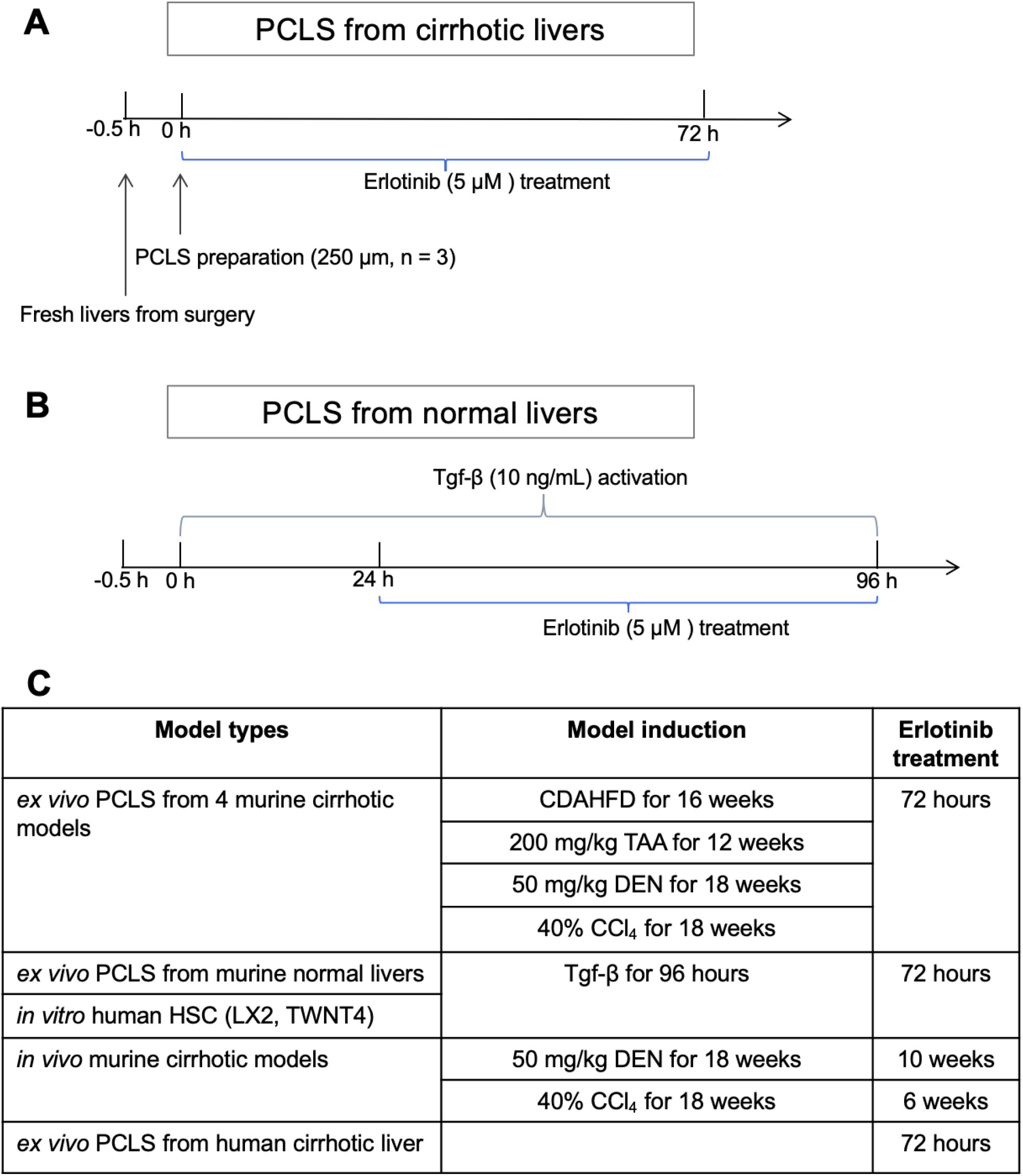
Schedule of PCLS preparation and erlotinib treatment. Workflow of PCLS preparation and erlotinib treatment for (A) cirrhotic and (B) normal livers. (C) Summary table of PCLS preparation and erlotinib treatment.

**Supplementary Figure 2.**
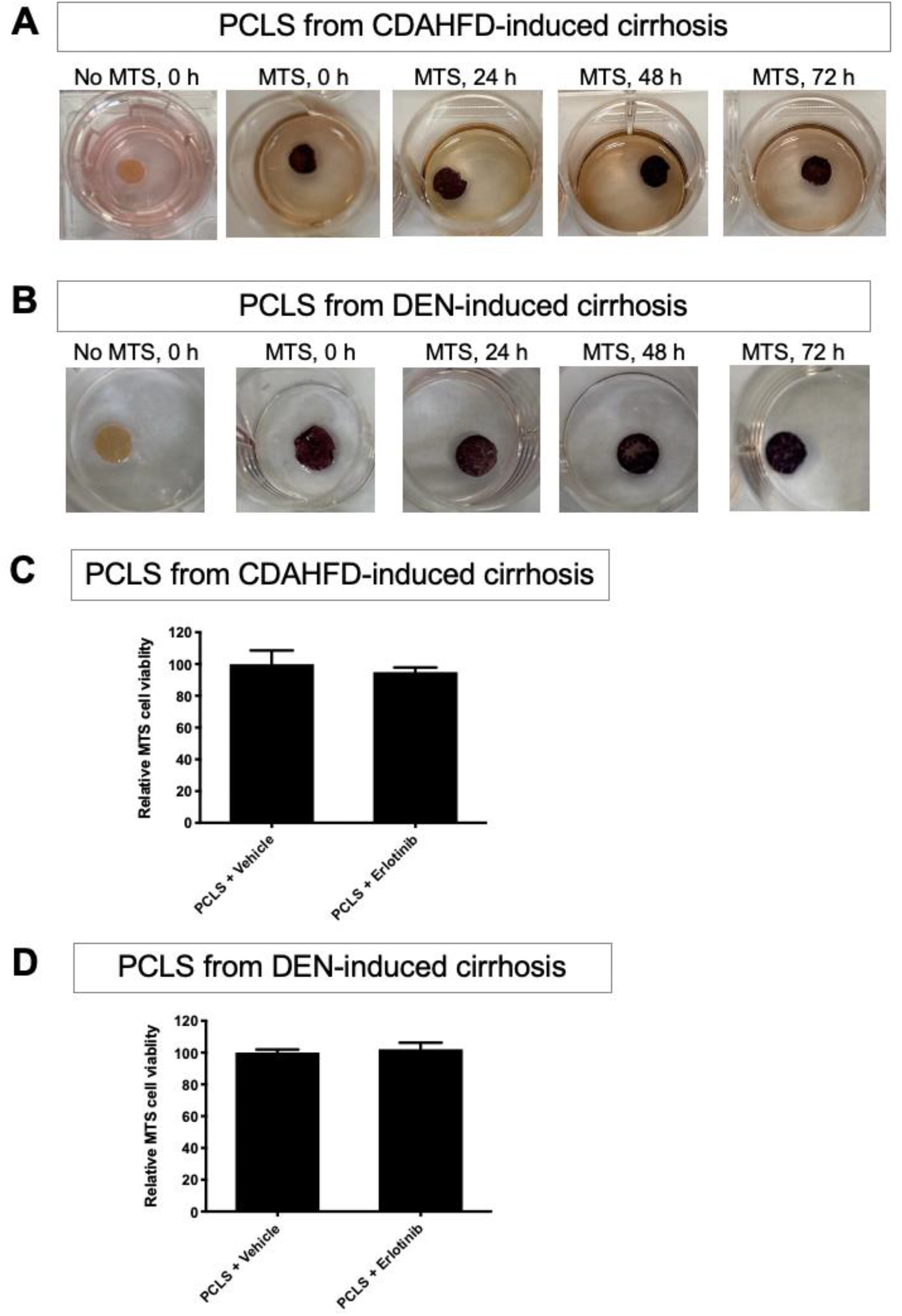
Viability evaluation in PCLS. (A) Representative images of PCLS from CDAHFD-induced cirrhosis rats on MTS assays, the cells alive were stained with purple color. (B) Representative images of PCLS from DEN-induced cirrhosis rats on MTS viability assays. MTS assays in PCLS from (C) CDAHFD and (D) DEN-induced rat cirrhosis after erlotinib treatment. These experiments were repeated three times (n = 3). All values were expressed as the mean ± S.E.M with two-tailed Student’s *t* tests.

**Supplementary Figure 3.**
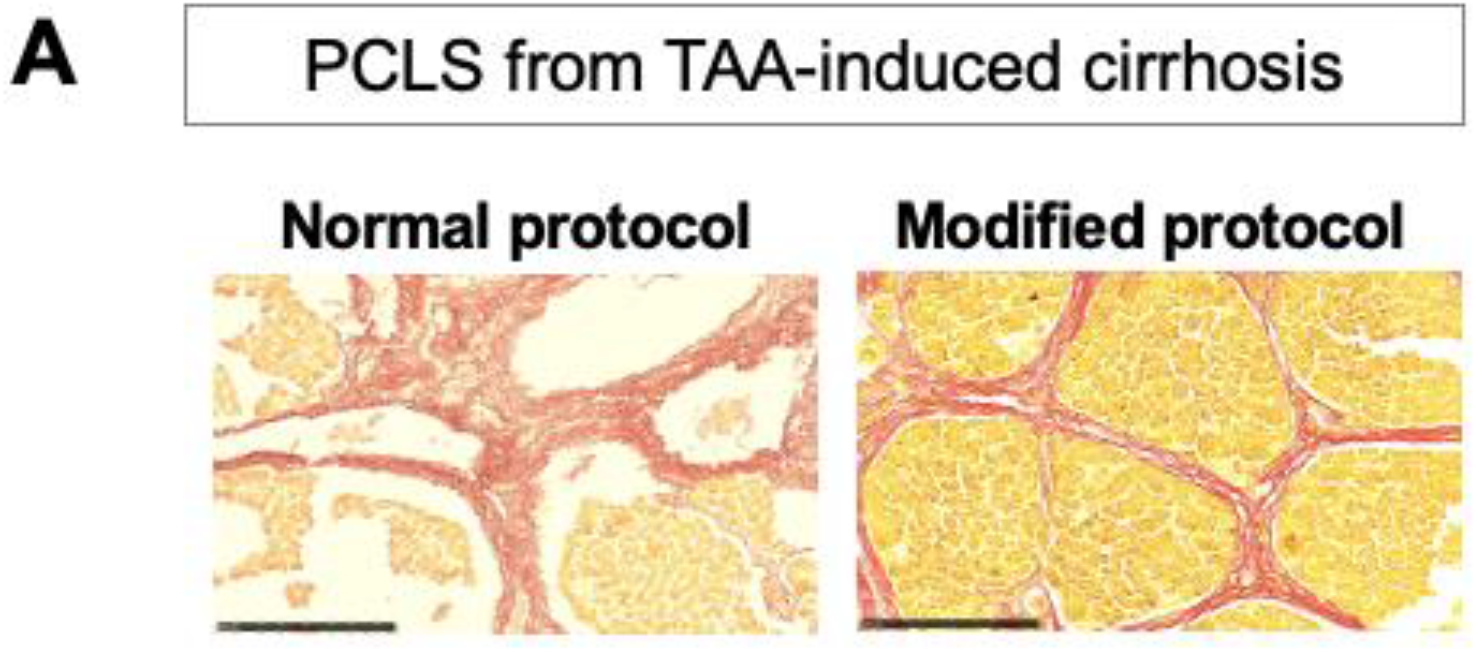
Improvement of PCLS processing. (A) Comparison of standard and modified protocols to proceed PCLS from TAA-induced rat cirrhosis.

**Supplementary Figure 4.**
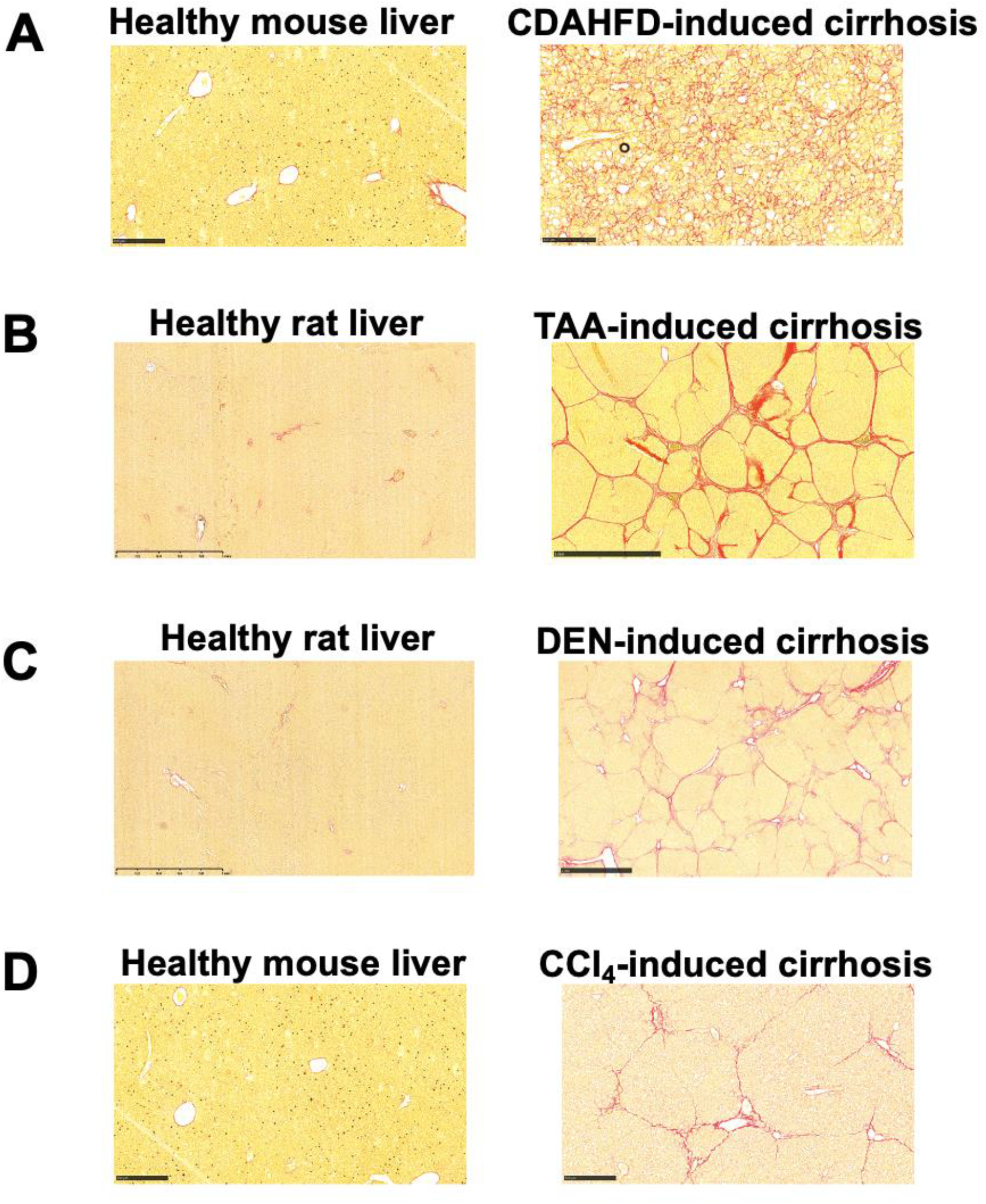
Confirmation of established murine cirrhosis. Confirmation of established (A) CDAHFD, (B) TAA, (C) DEN and (D) CCl_4_-induced murine cirrhosis.

**Supplementary Figure 5.**
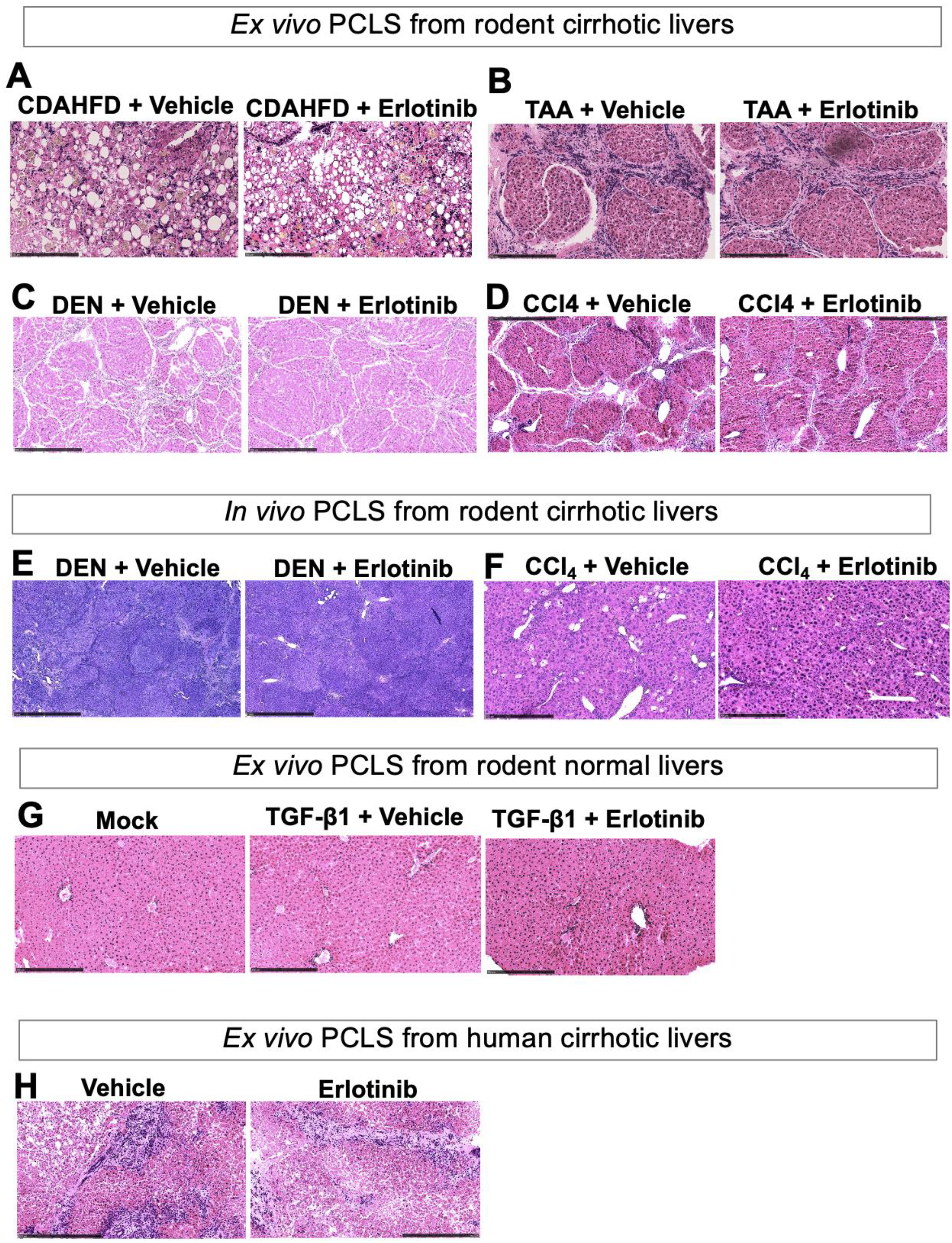
H&E staining. Representative images for H&E staining in PCLS from (A) CDAHFD-induced mouse cirrhosis, (B) TAA-induced rat cirrhosis, (C) DEN-induced *rat* cirrhosis, and (D) CCl_4_-induced *mouse* cirrhosis. (E) Representative images for H&E staining of PCLS from male healthy C57Bl/6 mice livers. (F) Representative images for H&E staining of human cirrhotic PCLS. Representative images for H&E staining of the samples from (G) *in vivo* DEN-induced rat cirrhotic models and (H) *in vivo* CCl_4_-induced mouse cirrhotic models.

**Supplementary Figure 6.**
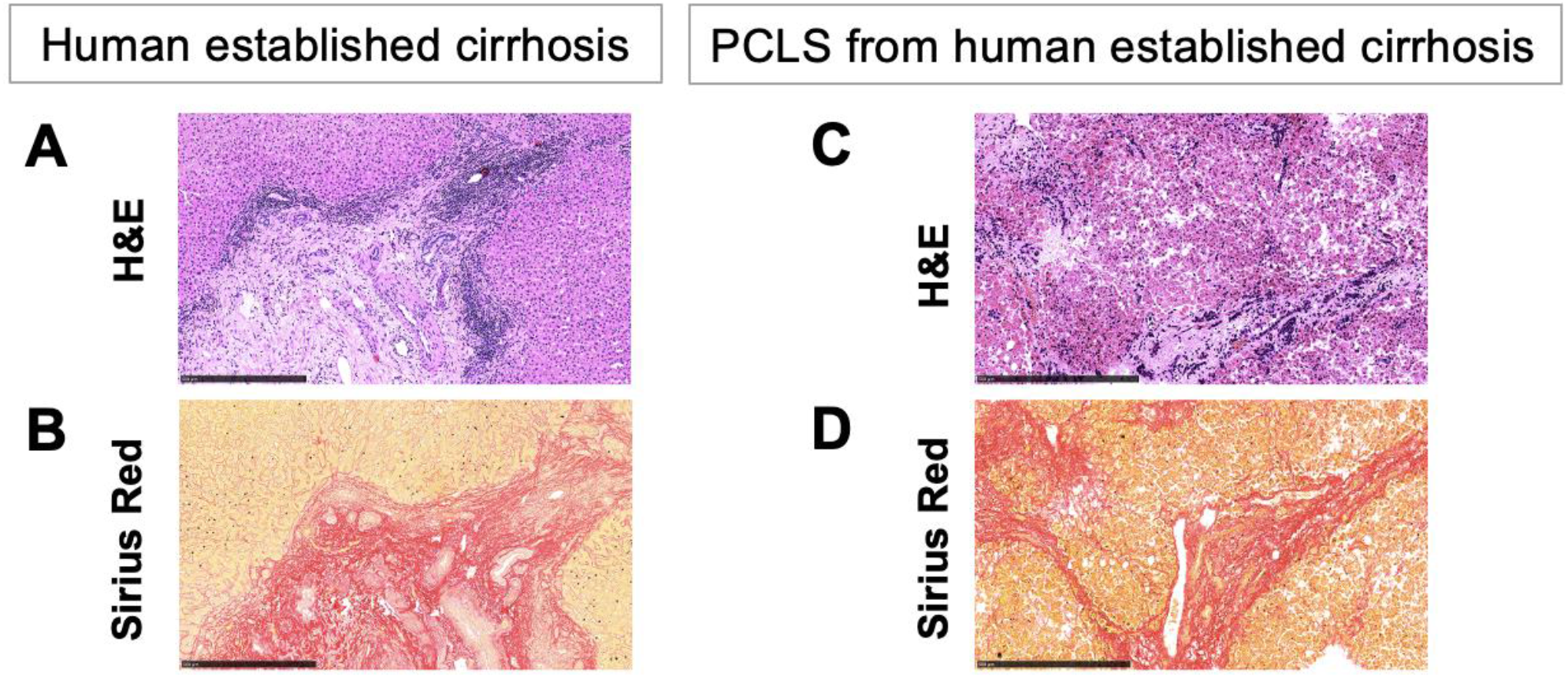
Confirmation of established human cirrhosis. Confirmation of human established cirrhosis with (A) H&E and (B) Sirius red staining. (C) H&E and (D) Sirius red staining of PCLS from human established cirrhosis.

**Supplementary Table 1.** Modified *Ishak* Scoring^31^.

| Score | Description |
| --- | --- |
| **0** | No fibrosis |
| **1** | Fibrous expansion of some or most portal areas, with or without short fibrous septa |
| **2** | Fibrous expansion of most portal areas, with occasional portal to portal bridging, or with marked portal to portal bridging as well as portal areas to central bridging |
| **3** | Marked bridging with occasional nodules |
| **4** | Probable to definite cirrhosis |

**Supplementary Table 2.**
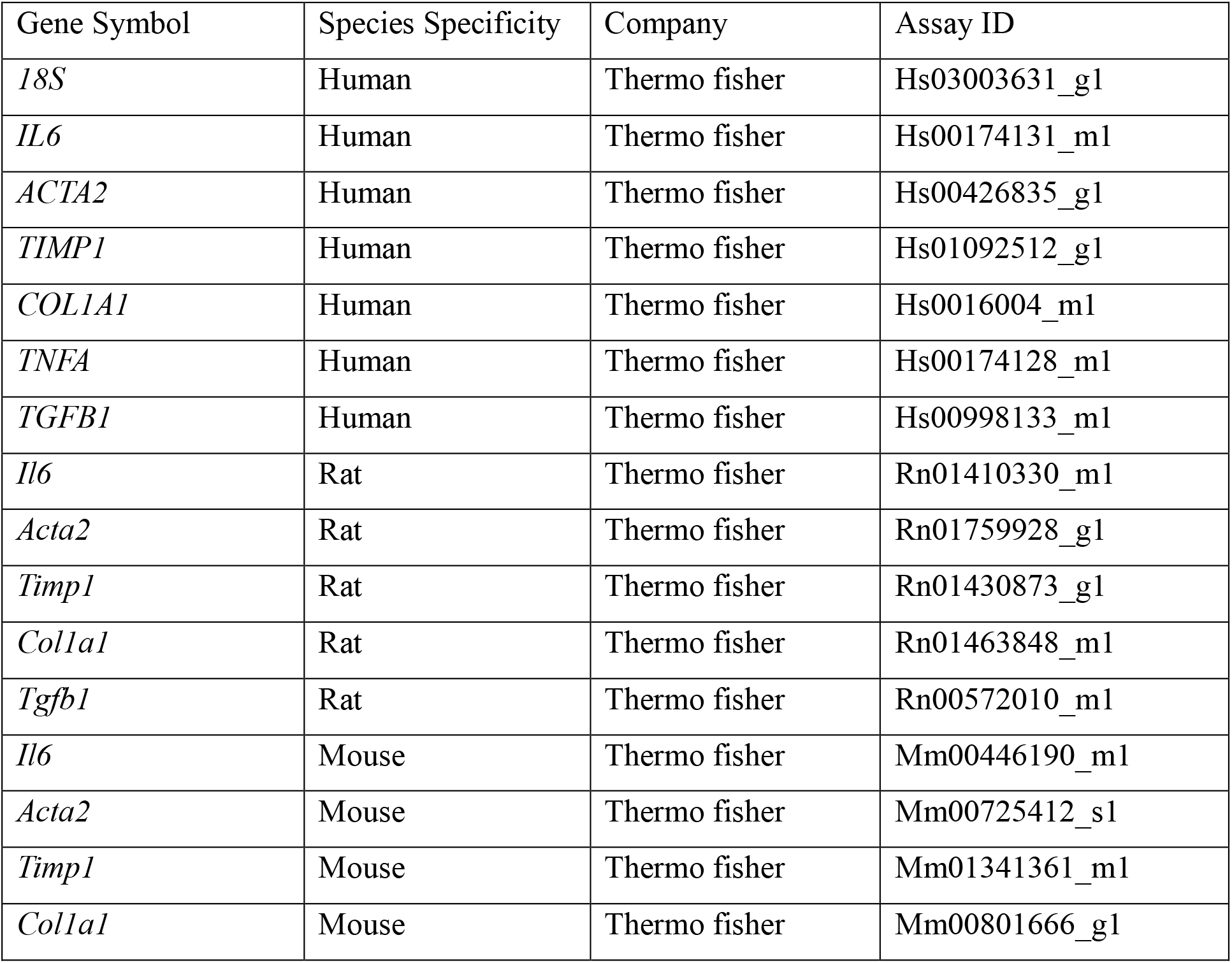
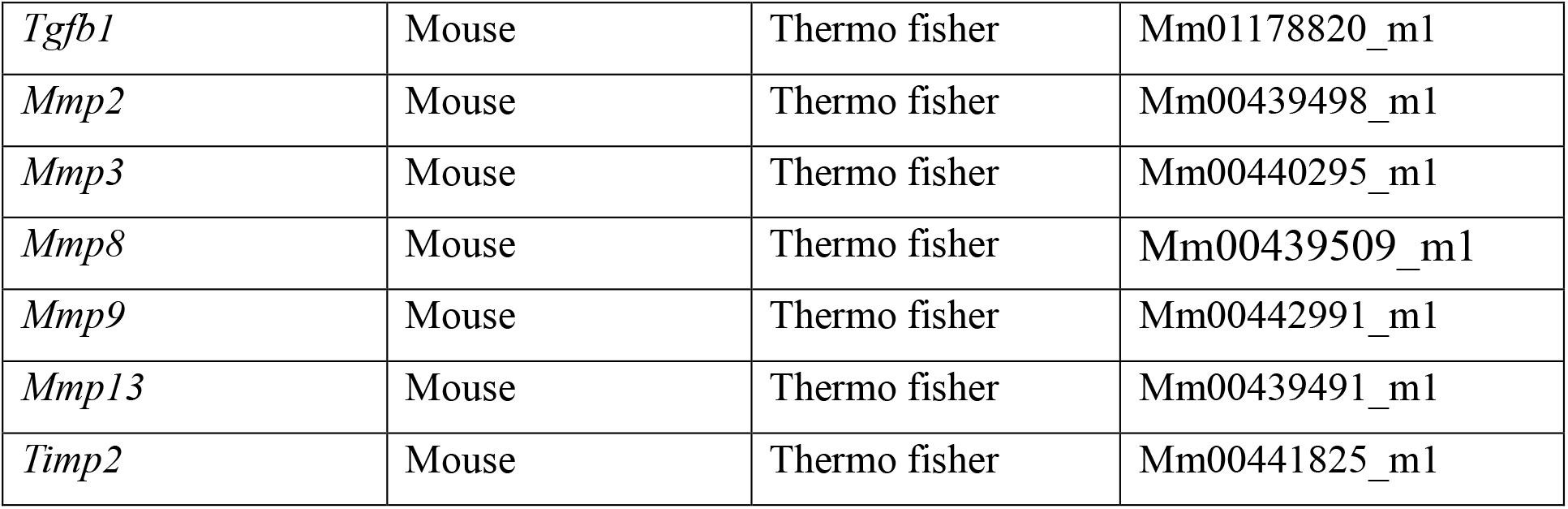
Oligonucleotides.

